# Gene expression signatures of salinity transitions in *Limia perugiae* (Poeciliidae), with comparisons to other teleosts

**DOI:** 10.1101/2023.12.21.572917

**Authors:** Elizabeth J. Wilson, Nick Barts, John Coffin, James B. Johnson, Carlos M. Rodríguez Peña, Joanna L. Kelley, Michael Tobler, Ryan Greenway

**Affiliations:** Division of Biology, Kansas State University, Manhattan, KS, USA; Department of Biology, University of Central Missouri, Warrensburg, MO, USA; Divison of Marine Fisheries, North Carolina Department of Environmental Quality, Morehead City, NC, USA; Instituto de Investigaciones Botánicas y Zoológicas, Universidad Autónoma de Santo Domingo, Santo Domingo, Dominican Republic; Department of Ecology and Evolutionary Biology, University of California Santa Cruz, Santa Cruz, CA, USA; Department of Biology, University of Missouri—St. Louis, St. Louis, MO 63121, USA; Whitney R. Harris World Ecology Center, University of Missouri—St. Louis, St. Louis, MO 63121, USA; WildCare Institute, Saint Louis Zoo, St. Louis, MO 63110, USA

**Keywords:** transcriptomics, poeciliid fishes, RNA-seq, convergence, osmoregulation, population divergence

## Abstract

Salinity gradients act as strong environmental barriers that limit the distribution of aquatic organisms. Changes in gene expression associated with transitions between freshwater and saltwater environments can provide insight into organismal responses to variation in salinity. We used RNA-sequencing (RNA-seq) to investigate genome-wide variation in gene expression between a hypersaline population and a freshwater population of the livebearing fish species *Limia perugiae* (Poeciliidae). Our analyses of gill gene expression revealed potential molecular mechanisms underlying salinity tolerance in this species, including the enrichment of genes involved in ion transport, maintenance of chemical homeostasis, and cell signaling in the hypersaline population. We also found differences in gene expression patterns associated with cell cycle and protein folding processes between the hypersaline and freshwater *L. perugiae*. Bidirectional freshwater-saltwater transitions have occurred repeatedly during the diversification of fishes, allowing for broad-scale examination of repeatable patterns in evolution. We compared transcriptomic variation in *L. perugiae* with other teleosts that have made freshwater-saltwater transitions to test for convergence in gene expression. Among the four distantly related population pairs from high- and low-salinity environments that we included in our analysis, we found only ten shared differentially expressed genes, indicating little evidence for convergence. However, we found that differentially expressed genes shared among three or more lineages were functionally enriched for ion transport and immune functioning. Overall, our results—in conjunction with other recent studies— suggest that different genes are involved in salinity transitions across disparate lineages of teleost fishes.

## Introduction

Variation in salinity imposes osmoregulatory challenges on aquatic organisms, and contact zones between freshwater and saltwater environments act as barriers that limit the ability of animals to move from one habitat to the other (Davis et al. 2012). Many aquatic taxa have consequently failed to cross natural salinity gradients (Lee & Bell 1999). Those that achieve such habitat shifts overcome osmoregulatory challenges through plasticity or adaptation, and both responses have greatly shaped aquatic species distributions (Corush 2019; Lee et al. 2003; Whitehead et al. 2011). Due to the ecological expansion that accompanies colonization of novel habitats, invasions from freshwater to saltwater environments, or vice versa, that result in adaptation are of particular interest for elucidating the genetic mechanisms underlying salinity tolerance and the diversification of aquatic organisms (Davis et al. 2012; Lee & Bell 1999; Betancur-R 2010).

Among fishes, diversification has coincided with repeated transitions between freshwater and saltwater habitats (Lee & Bell 1999). Many saltwater-freshwater transitions in fishes have occurred over long evolutionary timescales, and deep evolutionary divergences have resulted in many species only tolerating a narrow range of salinities, restricting them to either freshwater or saltwater environments (Betancur-R 2010; Lee & Bell 1999; Bloom et al. 2013; Bloom & Lovejoy 2017; Carrete Vega & Wiens 2012). A considerably smaller portion of fish species can survive in both freshwater and saltwater environments (Betancur-R et al. 2015). Among species that tolerate a broad range of salinities, movement along salinity clines is characteristic of diadromous lineages with life histories involving migration between freshwater and saltwater environments during an individual’s lifetime (Betancur-R et al. 2015; Bloom & Lovejoy 2014). While some species cannot cross the saltwater-freshwater boundary and some do so in their lifetimes, there are also lineages between these two extremes that have made transitions between saltwater and freshwater environments at microevolutionary scales (Betancur-R 2010; Kusakabe et al. 2017; Xu et al. 2013). Few studies have investigated mechanisms of salinity adaptation in species where closely related populations experience different salinity regimes, and the evolutionary repeatability of these mechanisms across lineages remains to be explored.

Transitions between freshwater and saltwater environments are challenging for animals that actively maintain internal solute homeostasis. The many physiological processes involved in stable state osmotic and ionic balance necessitate changes in multiple interdependent processes when crossing a salinity barrier (Kültz 2015). To maintain homeostasis, fishes in freshwater must actively absorb salt and excrete water in the form of dilute urine to counteract their passive loss of salt and absorption of water (Kültz 2015). In contrast, fishes in saltwater environments must remove salt and retain water (Kültz 2015). As a result, crossing a salinity barrier requires a shift between absorption and excretion of ions and water in multiple organs, involving both active and passive processes (Kültz 2015; Greenwell et al. 2003). Remodeling of gill epithelia and regulation of ion transporters, aquaporins, and tight junctions in the gill are particularly central to this process (Foskett et al. 1983; Greenwell et al. 2003; Hwang 1987; Velotta et al. 2017). One way to quantify such complex physiological responses to variation in salinity is to compare patterns of gene expression across populations in different habitat types.

Several studies have investigated the physiological and transcriptomic responses to changes in salinity between populations of the same species or among closely related species of fish (Gibbons et al. 2017; Hughes et al. 2017; Velotta et al. 2017; Xu et al. 2013). Differential gene expression associated with osmoregulation and ion transport is commonly found between populations inhabiting environments of different salinities (Hughes et al. 2017; Velotta et al. 2017; Xu et al. 2013). In addition, differential expression has also been documented in genes associated with other biological functions, including immune processes (Gibbons et al. 2017; Hughes et al. 2017), cell communication (Xu et al. 2013), stress tolerance (Xu et al. 2013), and gill membrane permeability (Gibbons et al. 2017). These studies highlight candidate pathways that play important roles in divergence between saltwater and freshwater ecotypes within species, but there have been few comparisons investigating the repeatability of gene expression responses to variation in salinity across phylogenetically disparate taxa. Investigating evidence of convergence in gene expression patterns across taxa that have undergone similar salinity transitions will provide insight into possible shared and unique pathways involved in adaptation to different salinity regimes.

In this study, we investigated patterns of gene expression in a freshwater and a hypersaline population of a livebearing fish species, *Limia perugiae* (Poeciliidae), and compared transcriptomic variation between these populations to those observed in other species that have undergone similar salinity transitions. Freshwater fishes in the genus *Limia* are endemic to the islands of the West Indies (Rauchenberger 1988; Weaver, Cruz, et al. 2016). *Limia perugiae* is a widespread species across the southern portion of Hispaniola, occurring in freshwater artesian springs and low-order creeks, as well as hypersaline inland lakes and coastal lagoons (Weaver, Tello, et al. 2016; Erbelding-Denk et al. 1994; Haney & Walsh 2003). Exposure to high salinities in *L. perugiae* has been shown to decrease metabolic rate (Haney & Walsh 2003), increase the production of Na^+^/K^+^-ATPase and oxidative phosphorylation proteins in the gills (Weaver, Tello, et al. 2016), and reduce adult body size (Weaver, Tello, et al. 2016). Though predominantly associated with freshwater habitats, many fishes in the family Poeciliidae are able to tolerate a broad range of salinities, a factor potentially responsible for facilitating their dispersal across a wide geographic range (Myers 1949; Smith & Bermingham 2005; Rosen & Bailey 1963). While the mechanisms and consequences of salinity tolerance at the biochemical and physiological levels have been a focus of research in poeciliids (Chervinski 1984; Gonzalez et al. 2005; Tsai et al. 2018; Weaver, Tello, et al. 2016; Yang et al. 2011), the genetic underpinnings of salinity tolerance have yet to be investigated in this family. We used a natural system with conspecific populations occurring in both a freshwater and hypersaline habitat to characterize the potential molecular mechanisms underlying high salinity tolerance in poeciliid fishes.

We used RNA-sequencing (RNA-seq) to study genome-wide patterns of gene expression between freshwater and hypersaline *L. perugiae*. From this analysis, we aimed to identify genes and physiological pathways associated with salinity tolerance in this species. We then compared the *L. perugiae* population pair to other population pairs of freshwater and saltwater ecotypes in disparate taxa to understand if mechanisms of osmoregulatory capability are shared across divergent lineages of teleost fishes. We utilized a comparative transcriptomics approach that leverages new and pre-existing gene expression datasets to address the following questions: 1) What genes are differentially expressed between freshwater and saltwater *L. perugiae* populations, and with what physiological processes are they associated? 2) Is there evidence for commonalities in gene expression among phylogenetically disparate teleosts with freshwater and saltwater populations?

## Materials and Methods

### Sample collection

*Limia perugiae* were collected using a seine from a hypersaline lagoon (Laguna Oviedo: 17.801 °N, 71.363 °W) and a geographically proximate freshwater stream (Los Cocos: 17.905 °N, 71.286 °W) in the Dominican Republic. Following capture, adult females (*N*=6 per site) were euthanized, and both sets of gills were extracted using sterilized scissors and forceps. Tissues were immediately preserved in RNAlater (Ambion Inc., Austin, TX, USA).

### RNA-seq library preparation

RNA extraction, library preparation, and sequencing of samples followed procedures previously employed for related poeciliid species (Kelley et al. 2012, 2016). Briefly, 10–30 mg of tissue from each individual was frozen in liquid nitrogen, pulverized, and total RNA was extracted using the NucleoSpin RNA kit (Machery-Nagel, Düren, Germany). mRNA isolation and cDNA library preparation were completed with the NEBNext Poly(A) mRNA Magnetic Isolation Module (New England Biolabs, Inc., Ipswich, MA, USA) and NEBNext Ultra Directional RNA Library Prep Kit for Illumina (New England Biolabs, Inc., Ipswich, MA, USA), with minor modifications to the manufacturers’ protocol (Kelley et al. 2012, 2016; Passow et al. 2017). cDNA libraries were individually barcoded, quantified with Qubit and an Agilent 2100 Bioanalyzer High Sensitivity DNA chip, and then pooled with cDNA samples from other projects in sets of 11–12 samples based on nanomolar concentrations. Samples were split across pools such that samples from each habitat type were not all sequenced together, and there was no evidence for lane effects. Libraries were sequenced on an Illumina HiSeq 2500 using paired-end 101-base-pair (bp) reads at the Washington State University Spokane Genomics Core.

### Mapping

All raw reads were trimmed twice (quality 0 to remove Illumina adapters, followed by quality 24) using Trimgalore! (v.0.4.0; Krueger 2014). Trimmed reads were mapped to the *Poecilia mexicana* reference genome (RefSeq accession number: GCF_001443325.1; Warren et al. 2018) with an appended mitochondrial genome (GenBank Accession Number: KC992998.1) using BWA-MEM v.0.7.12 (Li & Durbin 2009). We annotated genes from the *P. mexicana* reference genome by extracting the longest transcript for each gene (with the perl script gff2fast.pl: https://github.com/ISUgenomics/common_scripts/blob/master/gff2fasta.pl) and comparing them against entries in the human SWISS-PROT database (critical E-value 0.001; access date 04/15/2017) using BLASTx (Camacho et al. 2009). Each gene was annotated with the best BLAST hit from the human database based on the top high-scoring segment pair.

#### Differential gene expression

We used STRINGTIE (v.1.3.3b; Pertea et al. 2015, 2016) to quantify the number of reads mapped to each gene for each individual (measured in counts per million mapped reads) and then used the prepDE.py script (provided with STRINGTIE) to generate a read count matrix (Pertea et al. 2016). We removed genes that did not have at least two counts per million in three or more individuals across both populations, resulting in 18,659 genes that were included in differential gene expression analysis. We identified differentially expressed genes using generalized linear models (GLMs) in R, as implemented in the Bioconductor package edgeR (Robinson et al. 2010). We fit a negative binomial GLM to the normalized read counts of each gene based on tagwise dispersion estimates and a design matrix describing the comparison between the saltwater and freshwater population using glmFit. We assessed statistical significance using the GLM likelihood-ratio test with a false discovery rate (FDR) of q-value < 0.05, calculated with the Benjamini-Hochberg correction (Benjamini & Hochberg 1995). After identifying the set of differentially expressed genes between the saltwater and freshwater population, we used a Gene Ontology (GO) enrichment analysis to explore putative biological functions of these genes. We annotated all differentially expressed genes that had a match in the human SWISS-PROT database with GO IDs (Harris et al. 2004) and tested for the enrichment of specific GO IDs separately in up and downregulated genes relative to the full background set of 18,659 genes using GOrilla (FDR q-value < 0.05, accessed May 26, 2022; Eden et al. 2009). A total of 10,935 genes in the background set were associated with a GO term in the database.

### Weighted gene co-expression network analysis

We constructed weighted gene co-expression networks to identify clusters of genes that were co-expressed across our samples (Zhang & Horvath 2005). To prepare the gene expression data for this analysis, we applied a variance-stabilizing transformation to the filtered reads using the varianceStabilizingTransformation function from the DESeq2 package (v.1.36.0; Love et al. 2014) in R, which allowed us to normalize the read counts relative to library size and appropriately scale the data for clustering (Langfelder & Horvath 2008, 2012). After transforming and normalizing the read counts, we used the cpm function in the edgeR package (v.3.38.1; Robinson et al. 2010) to generate a gene matrix of scaled read counts (log_2_-cpm, counts per million mapped reads) from the transformed read counts (Coffin et al. 2022).

As outlined in the WGCNA package documentation (Langfelder & Horvath 2008), we used hierarchical clustering to cluster the samples based on their gene expression profiles. We used the hclust function from the flashClust package (v.1.1.2; Langfelder & Horvath 2012) to cluster the samples. We then created a weighted network adjacency matrix using the adjacency function from the WGCNA package (v.1.71; Langfelder & Horvath 2008, 2012). The adjacency matrix was constructed by calculating pairwise co-expression similarities (Pearson correlation coefficients) and raising them to a power of β, a soft thresholding power (Langfelder & Horvath 2008). We used the pickSoftThreshold function in the WGCNA package (Langfelder & Horvath 2008) to assist in selecting a value of β that ensured our network fit the approximate scale free topology criterion while retaining the highest possible mean connectivity between the network genes. Based on the scale free topology model fit and mean connectivity of our network, we selected β = 7.

From our correlation network, we then generated a topological overlap dissimilarity matrix to identify modules of co-expressed genes. We calculated dissimilarity between the genes by converting the adjacencies into topological overlap similarities using the TOMsimilarity function in the WGCNA package (Langfelder & Horvath 2008) and then subtracting these topological overlap measures from 1. To identify modules of co-expressed genes, we used the hclust function from the flashClust package (Langfelder & Horvath 2012) for hierarchical clustering of the genes and then created a hierarchical clustering dendrogram. We used the cutreeDynamic function from the dynamicTreeCut package (Langfelder & Horvath 2008; Langfelder et al. 2008) to extract the modules from the dendrogram. To summarize the gene expression variation in each module, the first principal component of each module in the expression matrix (the eigengene) was calculated using the moduleEigengenes function from the WGCNA package (Langfelder & Horvath 2008). We used the function mergeCloseModules in the WGCNA package to merge eigengenes that were highly correlated. We included eigengenes with a correlation greater than 0.9 for merging. Finally, we identified modules that were significantly associated with the presence or absence of salinity by calculating correlation coefficients between the eigengenes and the habitat type. *P*-values of the correlation coefficients were calculated using the corPvalueStudent function from the WGCNA package, and we retained modules with *P*-values less than 0.01 for functional enrichment analyses. Similar to our differential gene expression analysis, we used GOrilla (Eden et al. 2009) for Gene Ontology enrichment analysis of genes contained in modules exhibiting significant correlations with habitat type.

### Comparisons of L. perugiae with other species

We mined previously published datasets to identify gene expression patterns commonly associated with salinity tolerance in disparate taxa, including South American silversides (*Odontesthes bonariensis* and *Odontesthes argentinensis*) from Hughes et al. (2017), three-spine sticklebacks (*Gasterosteus aculeatus*) from Gibbons et al. (2017), Amur ide (*Leuciscus waleckii*) from Xu et al. (2013), and *L. perugiae* (this study; Table 1). Each of these experiments generated paired-end RNA-seq raw reads from gill tissue in two ecotypes (one freshwater and the other saltwater) of the same lineage.

**Table 1.** Species included in the analysis, including environment (freshwater [FW] or saltwater [SW]), collection location, sample size (N), NCBI Sequence Read Archive (SRA) accession numbers, and study reference.

| Species | Environment | Collection Location | N | SRA Accessions | Study Reference |
| --- | --- | --- | --- | --- | --- |
| *Limia perugiae* | FW | Los Cocos (17.905°N, 71.286°W), Dominican Republic | 6 | *Pending* | This study |
| *Limia perugiae* | SW | Laguna Oviedo (17.801°N, 71.363°W), Dominican Republic | 6 | *Pending* | This study |
| *Gasterosteus aculeatus* | FW | Trout Lake (49°30029"N, 123°52029"W), British Columbia, Canada | 5 | SRX2544970, SRX2544969, SRX2544968, SRX2544967, SRX2544966 | (Gibbons et al. 2017) |
| *Gasterosteus aculeatus* | SW | Oyster Lagoon (49°36043.53"N, 124°01052.12"W), British Columbia, Canada | 5 | SRX2544985, SRX2544984, SRX2544983, SRX2544982, SRX2544981 | (Gibbons et al. 2017) |
| *Odontesthes bonariensis* | FW | Lake Chascomús (35°34'S, 58°01'W), Argentina | 3 | SRX1681471, SRX1681473, SRX1681474 | (Hughes et al. 2017) |
| *Odontesthes argentinensis* | SW | Mar del Plata (38°02'S, 57°31'W), Argentina | 3 | SRX1671790, SRX1681012, SRX1681017 | (Hughes et al. 2017) |
| *Leuciscus waleckii* | FW | Ganggeng Nor Lake (43°17'48"N, 116°53'27"E), Mongolia | 1 | SRX333071 | (Xu et al. 2013) |
| *Leuciscus waleckii* | SW | Dali Nor Lake (43°22'43"N, 116°39'24"E), Mongolia | 1 | SRX1410650 | (Xu et al. 2013) |

Raw RNA-seq reads from each transcriptomics project were downloaded in FASTQ format, and reference genomes or transcriptomes for each species were downloaded in FASTA format from Genbank (see Table 1 for accession numbers). Reads were trimmed and mapped to their respective reference genomes (*Cyprinus carpio*: GCF_000951615.1, Xu et al. 2014; *Poecilia mexicana*: GCF_001443325.1, Warren et al. 2018; *Gasterosteus aculeatus*: Broad S1 v. 93, Jones et al. 2012) or reference transcriptome (*Menidia menidia*: GEVY00000000.1, Therkildsen & Palumbi 2017) following the same methods described above for *L. perugiae*. We then quantified the number of reads mapped to each gene in the annotation file for each reference genome and created a read counts matrix for each species, which were used for subsequent expression analyses. Expression analyses were performed in R version 4.1.2. The 10,000 genes with the highest standard deviation between freshwater and saltwater samples were abstracted from each read counts matrix, and overall expression patterns were visualized with multi-dimensional scaling (MDS) plots.

To make comparisons across species, we used OrthoFinder v2.2.6 to identify orthologous genes among the reference genomes (Emms & Kelly 2015, 2019), and 18,419 orthogroups were identified. To calculate counts per orthogroup, we used the gene counts matrix of each species to sum up the counts across all loci contained in an orthogroup. Based on this orthogroup counts matrix, we retained only the orthogroups that were expressed in all individuals (cpm > 0 per individual), resulting in 12,743 retained orthogroups.

To evaluate expression differences for each orthogroup, we made pairwise comparisons between ecotypes following the same methods described above for the *L. perugiae* comparisons. Briefly, we normalized reads, created and compared generalized linear models of the normalized read counts, generated a design matrix, estimated tagwise dispersion, and conducted GLM likelihood-ratio tests to test whether differences in expression were statistically different between the freshwater and saltwater population for each orthgroup. To identify orthogroups exhibiting convergent expression patterns across lineages, we intersected the significantly upregulated and downregulated orthogroups from all lineage-specific comparisons, identifying orthogroups that were differentially expressed in the same direction in pairwise, three-, and four-way comparisons among the lineages. After identifying the set of orthogroups with differential expression across three or more lineages, we used a GO enrichment analysis as described above to explore the putative biological functions of these candidate gene sets.

## Results

### Comparative analysis of freshwater and hypersaline L. perugiae

We used RNA-seq to characterize the transcriptomes of *L. perugiae* from a freshwater (n=6) and a hypersaline populations (Table 1). 73,590,325 total raw reads were obtained across all individuals: 34,998,157 from freshwater *L. perugiae* (n=6) and 38,592,168 from saltwater *L. perugiae* (n=6) before trimming (Table S1). After trimming, 95.5% of reads from the freshwater individuals mapped to the *Poecilia mexicana* reference genome, and 95.2% mapped for the saltwater individuals (Table S1).

We identified 4,895 differentially expressed genes between saltwater and freshwater ecotypes of *L. perugiae*, 2,437 of which were upregulated and 2,458 of which were downregulated in the saltwater ecotype (Figure 1A). The genes upregulated in the saltwater population were largely associated with ion transport, maintaining chemical homeostasis, and cell signaling (FDR < 0.05) (Table 2). Processes relevant to chemical homeostasis included several solute carrier genes, such as *SLC9A3*, *SLC8B1*, *SLC12A8*, and *SLC30A9*. Na^+^/K^+^-ATPase and other ATPase genes, such as *ATP1B1* and *ATP6V1A*, were also upregulated among the genes involved in chemical homeostasis and signal transduction. Genes downregulated in the saltwater population corresponded to GO process terms such as mitotic cell cycle process, protein folding, chromosome segregation, rRNA processing, and mitochondrial translational elongation (FDR < 0.05; Table 2).

**Figure 1.**
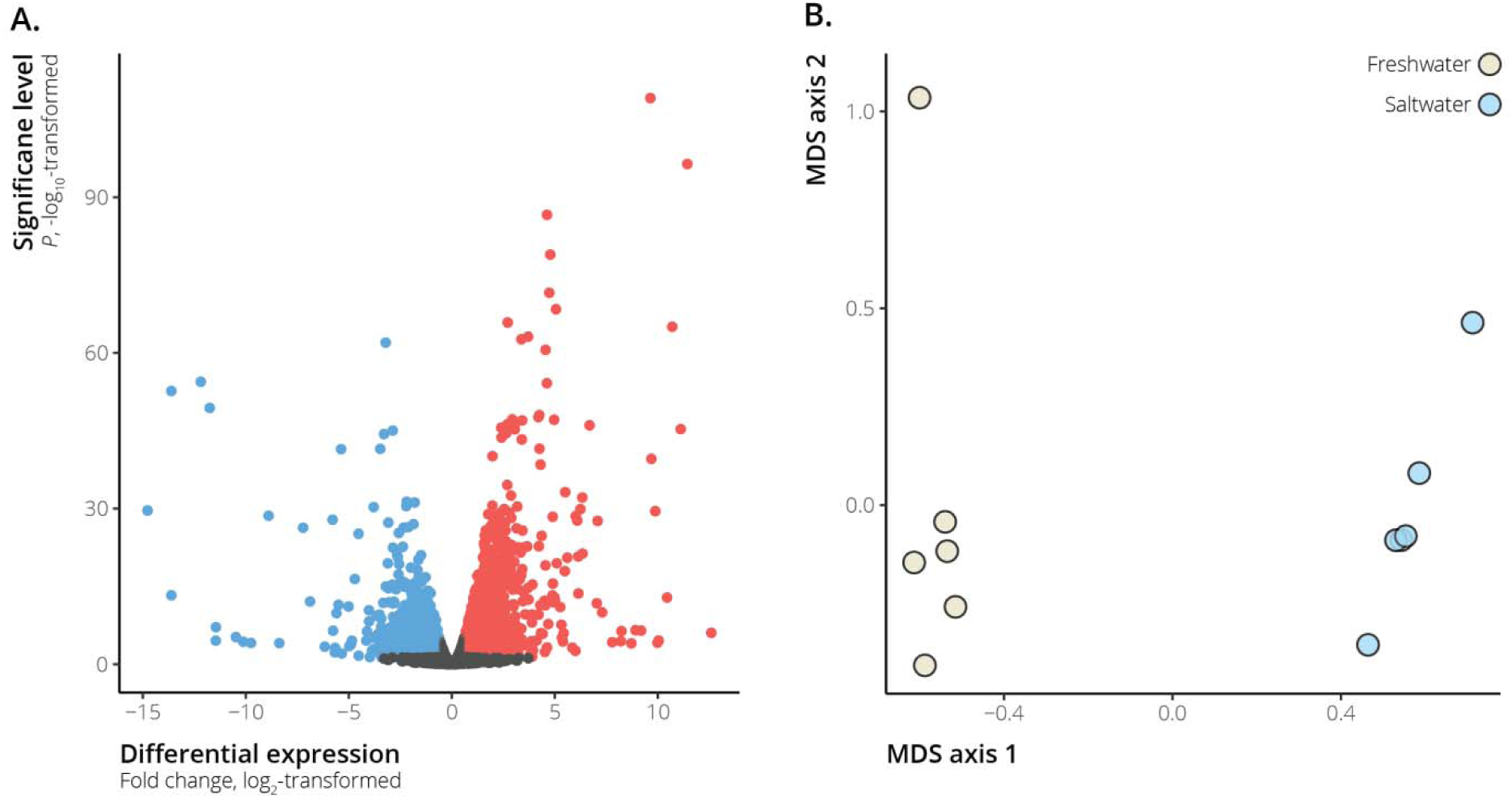
A. Volcano plot depicting differentially expressed genes between hypersaline and freshwater *Limia perugiae*. Genes that were significantly differentially expressed between hypersaline and freshwater populations (FDR < 0.05) are indicated by the blue and red points—blue points represent genes downregulated in the hypersaline populations, while red points represent upregulated genes. B. Multi-dimensional scaling (MDS) plot of hypersaline and freshwater *L. perugiae* gene expression profiles. MDS axis 1 separated samples by freshwater vs. hypersaline environments.

**Table 2.** GO process terms with significant enrichment in genes upregulated and downregulated in the hypersaline ecotype of *Limia perugiae* (FDR < 0.05). The table includes the GO term ID, description, the degree of enrichment, the number of differentially expressed genes associated with the GO term (*N*), as well as *P* and FDR-corrected *q*-values.

| GO term | Description | Enrichment | *N* | *P*-value | FDR *q*-value |
| --- | --- | --- | --- | --- | --- |
| **Upregulated** |  |  |  |  |  |
| GO:0006820 | anion transport | 1.8 | 100 | 5.67E-10 | 8.04E-06 |
| GO:0006811 | ion transport | 1.49 | 181 | 3.41E-09 | 2.42E-05 |
| GO:0007165 | signal transduction | 1.22 | 540 | 4.67E-09 | 2.21E-05 |
| GO:0051049 | regulation of transport | 1.36 | 242 | 4.50E-08 | 1.59E-04 |
| GO:0034220 | ion transmembrane transport | 1.51 | 129 | 3.41E-07 | 9.67E-04 |
| GO:0043269 | regulation of ion transport | 1.61 | 98 | 3.89E-07 | 9.19E-04 |
| GO:0010959 | regulation of metal ion transport | 1.85 | 62 | 4.53E-07 | 9.18E-04 |
| GO:0098656 | anion transmembrane transport | 1.89 | 57 | 5.26E-07 | 9.33E-04 |
| GO:0032879 | regulation of localization | 1.24 | 360 | 1.19E-06 | 1.87E-03 |
| GO:0015711 | organic anion transport | 1.71 | 73 | 1.21E-06 | 1.71E-03 |
| GO:0002028 | regulation of sodium ion transport | 2.72 | 23 | 1.62E-06 | 2.09E-03 |
| GO:0023051 | regulation of signaling | 1.19 | 451 | 3.07E-06 | 3.62E-03 |
| GO:0048878 | chemical homeostasis | 1.4 | 153 | 3.61E-06 | 3.94E-03 |
| GO:0010646 | regulation of cell communication | 1.19 | 444 | 7.08E-06 | 7.17E-03 |
| GO:0003254 | regulation of membrane depolarization | 3.24 | 15 | 7.84E-06 | 7.41E-03 |
| GO:0051050 | positive regulation of transport | 1.41 | 132 | 1.38E-05 | 1.23E-02 |
| GO:1902305 | regulation of sodium ion transmembrane transport | 2.85 | 17 | 1.76E-05 | 1.46E-02 |
| GO:0050794 | regulation of cellular process | 1.07 | 1144 | 2.50E-05 | 1.97E-02 |
| GO:0031344 | regulation of cell projection organization | 1.43 | 111 | 3.54E-05 | 2.64E-02 |
| GO:0043270 | positive regulation of ion transport | 1.78 | 46 | 3.61E-05 | 2.56E-02 |
| GO:0007186 | G protein-coupled receptor signaling pathway | 1.47 | 97 | 3.72E-05 | 2.51E-02 |
| GO:0010975 | regulation of neuron projection development | 1.51 | 84 | 3.89E-05 | 2.51E-02 |
| GO:0045664 | regulation of neuron differentiation | 1.44 | 105 | 4.04E-05 | 2.49E-02 |
| GO:0042908 | xenobiotic transport | 3.3 | 12 | 5.15E-05 | 3.04E-02 |
| GO:0019932 | second-messenger-mediated signaling | 1.69 | 53 | 5.17E-05 | 2.93E-02 |
| GO:0042592 | homeostatic process | 1.29 | 197 | 5.18E-05 | 2.83E-02 |
| GO:0120035 | regulation of plasma membrane bounded cell projection organization | 1.42 | 109 | 5.79E-05 | 3.04E-02 |
| GO:0006814 | sodium ion transport | 2.04 | 29 | 7.33E-05 | 3.71E-02 |
| GO:0032501 | multicellular organismal process | 1.19 | 360 | 7.62E-05 | 3.72E-02 |
| GO:0045216 | cell-cell junction organization | 1.94 | 33 | 7.65E-05 | 3.62E-02 |
| GO:0050767 | regulation of neurogenesis | 1.38 | 122 | 7.78E-05 | 3.56E-02 |
| GO:0055085 | transmembrane transport | 1.3 | 165 | 9.84E-05 | 4.36E-02 |
| GO:0030001 | metal ion transport | 1.47 | 85 | 1.05E-04 | 4.52E-02 |
| GO:0031345 | negative regulation of cell projection organization | 1.83 | 37 | 1.07E-04 | 4.46E-02 |
| GO:0065008 | regulation of biological quality | 1.15 | 453 | 1.21E-04 | 4.91E-02 |
| GO:0015849 | organic acid transport | 1.69 | 47 | 1.25E-04 | 4.91E-02 |
| GO:0046942 | carboxylic acid transport | 1.69 | 47 | 1.25E-04 | 4.77E-02 |
| GO:0065007 | biological regulation | 1.06 | 1274 | 1.27E-04 | 4.75E-02 |
| **Downregulated** |  |  |  |  |  |
| GO:1903047 | mitotic cell cycle process | 1.66 | 149 | 2.52E-11 | 3.58E-07 |
| GO:0006457 | protein folding | 2.14 | 64 | 3.03E-10 | 2.15E-06 |
| GO:0022402 | cell cycle process | 1.46 | 191 | 6.63E-09 | 3.13E-05 |
| GO:0051301 | cell division | 1.71 | 98 | 1.40E-08 | 4.98E-05 |
| GO:0006364 | rRNA processing | 1.91 | 65 | 4.51E-08 | 1.28E-04 |
| GO:0009987 | cellular process | 1.05 | 1687 | 6.13E-08 | 1.45E-04 |
| GO:0016072 | rRNA metabolic process | 1.82 | 69 | 1.35E-07 | 2.73E-04 |
| GO:0030198 | extracellular matrix organization | 1.75 | 76 | 2.13E-07 | 3.77E-04 |
| GO:0007059 | chromosome segregation | 2.38 | 33 | 3.54E-07 | 5.57E-04 |
| GO:0043062 | extracellular structure organization | 1.7 | 80 | 3.85E-07 | 5.46E-04 |
| GO:0000070 | mitotic sister chromatid segregation | 3.61 | 15 | 7.56E-07 | 9.74E-04 |
| GO:0000819 | sister chromatid segregation | 3.41 | 16 | 1.01E-06 | 1.19E-03 |
| GO:0007051 | spindle organization | 2.04 | 42 | 1.40E-06 | 1.53E-03 |
| GO:0043933 | protein-containing complex subunit organization | 1.28 | 269 | 2.07E-06 | 2.10E-03 |
| GO:0098813 | nuclear chromosome segregation | 3.14 | 17 | 2.34E-06 | 2.21E-03 |
| GO:0006415 | translational termination | 2.16 | 35 | 2.50E-06 | 2.22E-03 |
| GO:0043624 | cellular protein complex disassembly | 1.99 | 42 | 3.07E-06 | 2.56E-03 |
| GO:0070125 | mitochondrial translational elongation | 2.18 | 33 | 3.71E-06 | 2.92E-03 |
| GO:0070098 | chemokine-mediated signaling pathway | 3.17 | 16 | 4.01E-06 | 2.99E-03 |
| GO:0018208 | peptidyl-proline modification | 2.65 | 21 | 6.13E-06 | 4.34E-03 |
| GO:0007010 | cytoskeleton organization | 1.38 | 153 | 8.67E-06 | 5.85E-03 |
| GO:0007088 | regulation of mitotic nuclear division | 1.89 | 42 | 1.29E-05 | 8.34E-03 |
| GO:0044772 | mitotic cell cycle phase transition | 1.66 | 62 | 1.72E-05 | 1.06E-02 |
| GO:0006336 | DNA replication-independent nucleosome assembly | 3.1 | 14 | 2.24E-05 | 1.32E-02 |
| GO:0034622 | cellular protein-containing complex assembly | 1.36 | 150 | 2.47E-05 | 1.40E-02 |
| GO:0051783 | regulation of nuclear division | 1.82 | 44 | 2.48E-05 | 1.35E-02 |
| GO:0051383 | kinetochore organization | 4.16 | 9 | 2.59E-05 | 1.36E-02 |
| GO:0044770 | cell cycle phase transition | 1.63 | 63 | 2.67E-05 | 1.35E-02 |
| GO:0000278 | mitotic cell cycle | 1.94 | 36 | 2.98E-05 | 1.46E-02 |
| GO:0000413 | protein peptidyl-prolyl isomerization | 2.77 | 16 | 3.82E-05 | 1.80E-02 |
| GO:0034724 | DNA replication-independent nucleosome organization | 2.98 | 14 | 4.05E-05 | 1.85E-02 |
| GO:0070126 | mitochondrial translational termination | 2.02 | 31 | 4.10E-05 | 1.82E-02 |
| GO:0051276 | chromosome organization | 1.53 | 75 | 5.71E-05 | 2.45E-02 |
| GO:0071840 | cellular component organization or biogenesis | 1.11 | 739 | 6.92E-05 | 2.88E-02 |
| GO:0006414 | translational elongation | 1.85 | 37 | 7.46E-05 | 3.02E-02 |
| GO:0022610 | biological adhesion | 1.35 | 134 | 8.31E-05 | 3.27E-02 |
| GO:0007155 | cell adhesion | 1.35 | 133 | 8.91E-05 | 3.41E-02 |
| GO:0042493 | response to drug | 1.37 | 119 | 9.53E-05 | 3.56E-02 |
| GO:0016074 | snoRNA metabolic process | 3.46 | 10 | 9.90E-05 | 3.60E-02 |
| GO:0000281 | mitotic cytokinesis | 2.43 | 18 | 1.11E-04 | 3.95E-02 |
| GO:0000086 | G2/M transition of mitotic cell cycle | 1.83 | 36 | 1.15E-04 | 3.97E-02 |
| GO:1902850 | microtubule cytoskeleton organization involved in mitosis | 1.89 | 33 | 1.15E-04 | 3.88E-02 |
| GO:0045943 | positive regulation of transcription by RNA polymerase I | 3.21 | 11 | 1.16E-04 | 3.84E-02 |
| GO:0006334 | nucleosome assembly | 2.24 | 21 | 1.33E-04 | 4.30E-02 |
| GO:0044839 | cell cycle G2/M phase transition | 1.81 | 36 | 1.42E-04 | 4.46E-02 |
| GO:0072321 | chaperone-mediated protein transport | 4.31 | 7 | 1.58E-04 | 4.86E-02 |

WGCNA reveaked five modules of co-expressed genes that were significantly correlated with salinity (Figure 2). The turquoise module was positively correlated with salinity (*P*-value < 0.01), and the black, red, royalblue, and blue modules were all negatively correlated with salinity (Figure 2). Out of the 18,659 genes included in our analysis, the positively correlated module (turquoise module) contained 4,056 genes. The negatively correlated modules contained 2,166 genes (black module), 837 genes (red module), 173 genes (royalblue module), and 3,300 genes (blue module). The correlation coefficients and their associated *P*-values between each gene and the environmental condition (salinity), and between each gene and each module, are included in Table S2. Each gene’s module assignment can also be found in Table S2. From the functional enrichment analysis, we found that the turquoise, royalblue, and blue modules were significantly enriched for biological processes, and these modules of co-expressed genes largely corroborated the differential expression results. Like the biological processes that were enriched among the up-regulated genes in the saltwater population, the module that was positively correlated with salinity (turquoise) was functionally enriched for GO terms involved in ion transport and cell signaling (Table S3). There were, however, GO terms associated with the turquoise module that were not identified from the differential expression analysis, including regulation of autophagy and lipid transport (Table S3). The modules that were negatively correlated with salinity also reflected biological processes that were associated with down-regulated genes in the saltwater *L. perugiae*, including regulation of the cell cycle and protein folding (Table S4 and Table S5). The royalblue module only had one significantly enriched GO term, chemokine-mediated signaling pathway, which was a GO term also associated with down-regulation of genes in the saltwater environment (Table S4). In contrast, the blue module contained several significant GO terms, including mitotic cell cycle process, protein folding, rRNA metabolic process, and chromosome organization (Table S5). There were many GO terms associated with the blue module that were not represented in the differential expression analysis, including sarcomere organization, macromolecule biosynthetic process, regulation of cellular response to heat, and nucleic acid metabolic process (Table S5). Several of the GO terms unique to the blue module were related to cellular metabolism and biosynthesis pathways (Table S5).

**Figure 2.**
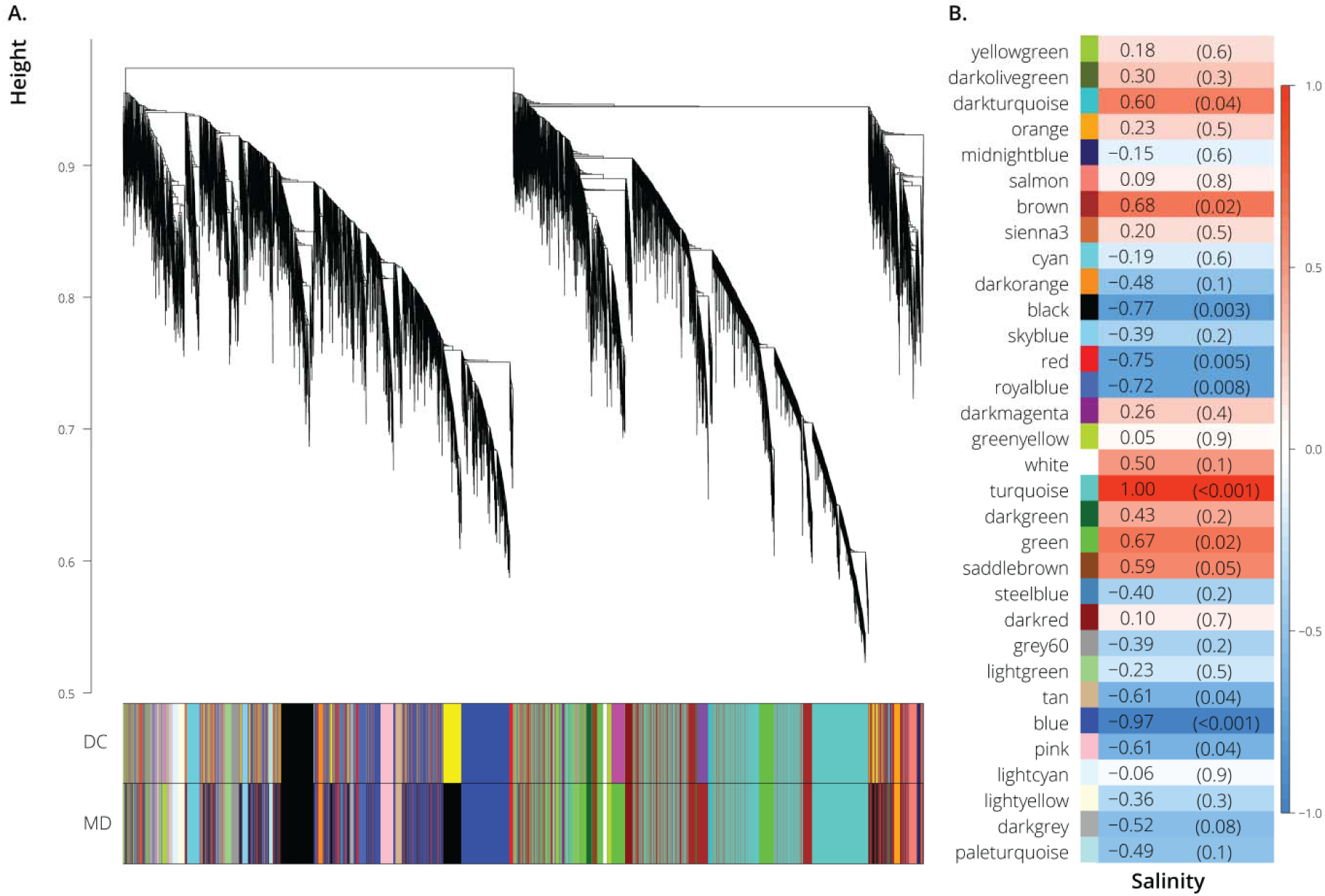
Weighted gene co-expression network analysis. A. Average linkage clustering tree based on topological overlap distances in gene expression patterns of *L. perugiae* from freshwater and saltwater habitats. Branches of the dendrogram correspond to modules, as shown in the color bars below. B. Correlation between module eigenvalues and habitat type (freshwater *vs*. saltwater). Each row corresponds to a module of coexpressed genes, and values are Pearson correlation coefficients (left column) and *P*-values (right column in parentheses). Color coloration scales with the correlation coefficient according to the scale bar to the right.

### Comparisons of Limia with phylogenetically disparate teleosts

We compared transcriptomic differences between *L. perugiae* populations to previously published transcriptome data from teleosts with populations from freshwater and saline habitats (Table 1). Mapping statistics for all four population pairs can be found in Supplementary Table 1. As expected, MDS plots indicated that orthogroup expression variation was primarily driven by differences among taxonomic groups, with much smaller differences between ecotypes within species (Figure 3A). We then compared orthogroup expression profiles of all the freshwater and saltwater lineages based on mean expression values and found that the variation in orthogroup expression largely reflects phylogenetic divergence among lineages (Figure 3B). Closely related lineages exhibited more similar expression profiles, irrespective of environmental conditions (saltwater vs. freshwater; Figure 3B). Across all groups, 122 differentially expressed orthogroups were shared across at least three lineages, but only 10 shared orthogroups were differentially expressed across all freshwater and saltwater population pairs (Figure 3C). Of those 10 orthogroups, 9 had annotations in the SWISS-PROT database (Table 3; Figure 4), including a Na^+^/H^+^-exchanger (*SLC9A3*) involved in osmoregulation. The directionality of differential expression varied among lineages, and none of the 10 shared differentially expressed orthogroups were up-regulated or down-regulated among all four population pairs (Figure 4).

**Figure 3.**
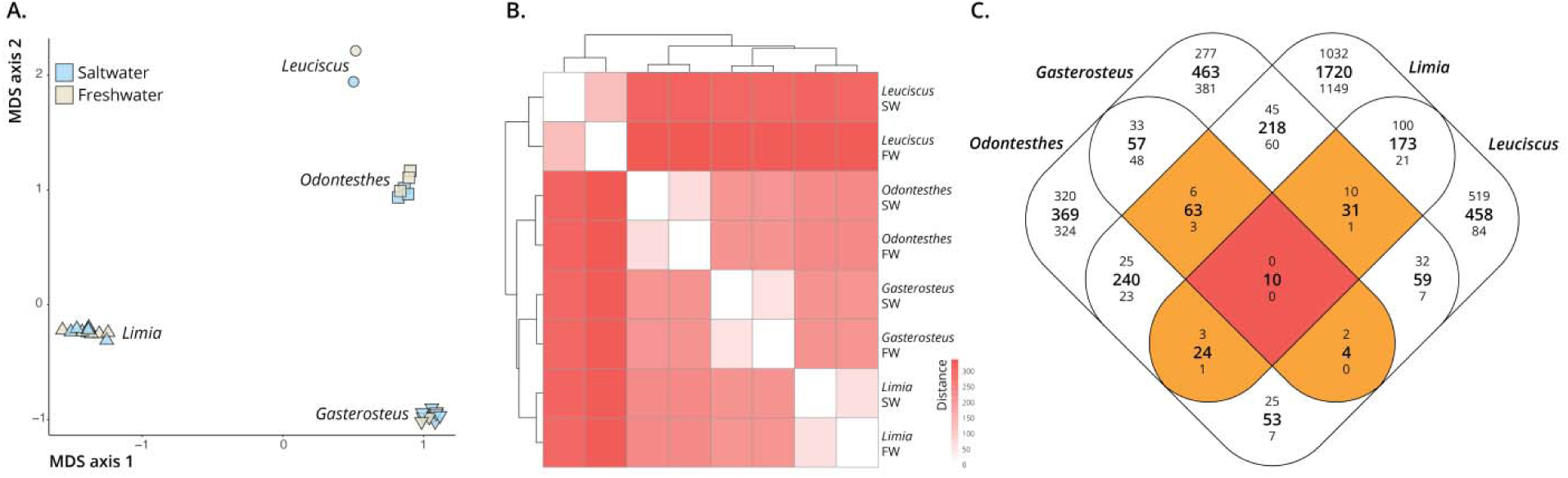
A. Multi-dimensional scaling plot (MDS) of the general expression patterns of all the populations included in our analysis. B. Similarity of gene expression profiles of saltwater (SW) and freshwater (FW) populations across different lineages. The majority of variation in gene expression reflects phylogenetic divergence among lineages. C. Shared differentially expressed genes across lineages. The large, central number in each section represents the total number of shared differentially expressed genes among the lineages in that intersection. The top number in each section represents the number of shared up-regulated genes, and the bottom number is the number of shared down-regulated genes in that intersection. Only 10 genes were consistently differentially expressed between all of the SW and FW populations.

**Figure 4.**
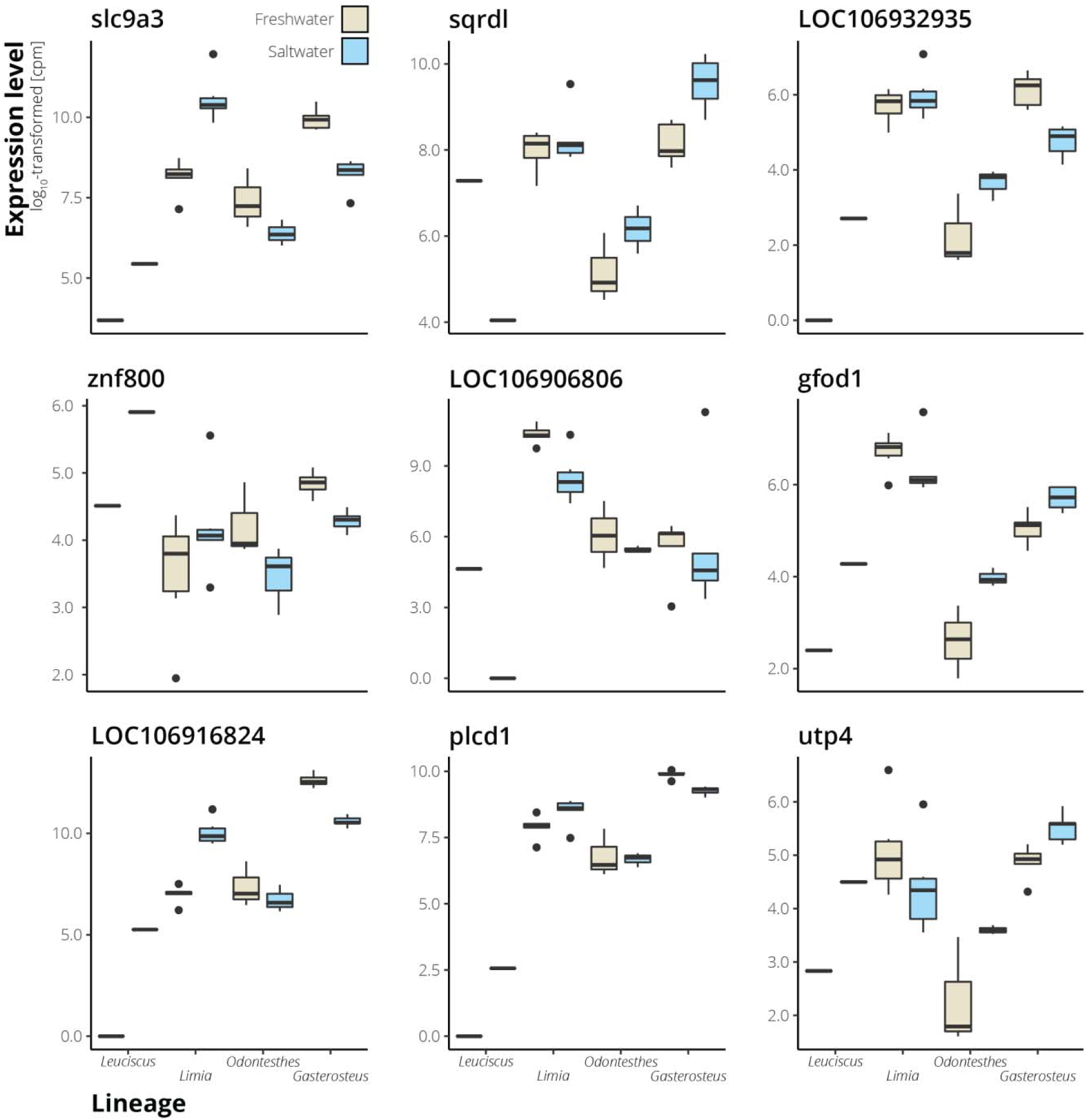
Examples of expression variation in shared differentially expressed genes. Nine of the ten shared differentially expressed genes have annotations, and those genes are included here. Magnitude and direction of differential expression is lineage specific.

**Table 3.** Shared differentially expressed genes among all four population pairs and their associated proteins. Based on the SWISS-PROT database, nine of the ten genes have experimental evidence for the existence of the protein associated with each gene. The tenth gene did not have a name match in the database, so it is not included.

| Protein name | Gene name |
| --- | --- |
| Sodium/hydrogen exchanger 3 | *SLC9A3* |
| Sulfide:quinone oxidoreductase, mitochondrial | *SQRDL* |
| Cytochrome b reductase 1 | *CYBRD1* |
| Zinc finger protein 800 | *ZNF800* |
| Pulmonary surfactant-associated protein D | *SFTPD* |
| Glucose-fructose oxidoreductase domain-containing protein 1 | *GFOD1* |
| Carcinoembryonic antigen-related cell adhesion molecule 1 | *CEACAM1* |
| 1-phosphatidylinositol 4,5-bisphosphate phosphodiesterase delta-1 | *PLCD1* |
| Cirhin | *CIRH1A* |

To investigate the biological processes reflected in shared differentially expressed orthogroups, we analyzed the differentially expressed orthogroups that were shared among three or more lineages (Figure 3C). GO analysis indicated enrichment in biological processes associated mostly with transmembrane transport (particularly ion transport) and some associated with immune function, which were all significant based on *P*-value (*P*-value < 0.001) but not after FDR correction (Table 4). Some of the GO processes specific to transmembrane transport included anion transport, inorganic cation import across plasma membrane, potassium ion import, urea transmembrane transport, and pyruvate transmembrane transport. The shared differentially expressed immune genes were reflective of negative regulation of T-helper cell differentiation, negative regulation of leukocyte differentiation, negative regulation of CD4-positive, alpha-beta T cell differentiation, and regulation of adaptive immune response based on somatic recombination of immune receptors built from immunoglobulin superfamily.

**Table 4.** GO process terms with significant enrichment in orthogenes with significant differential expression in at least three lineages (FDR < 0.05). The table include the GO term ID, description, the degree of enrichment, the number of differentially expressed genes associated with the GO term (*N*), as well as *P* and FDR-corrected *q*-values.

| GO term | Description | Enrichment | *N* | *P*-value | FDR *q*-value |
| --- | --- | --- | --- | --- | --- |
| GO:0006820 | anion transport | 3.63 | 14 | 2.81E-05 | 3.71E-01 |
| GO:0098656 | anion transmembrane transport | 4.99 | 10 | 2.90E-05 | 1.91E-01 |
| GO:0099587 | inorganic ion import across plasma membrane | 10.07 | 5 | 1.17E-04 | 5.16E-01 |
| GO:0098659 | inorganic cation import across plasma membrane | 10.07 | 5 | 1.17E-04 | 3.87E-01 |
| GO:0010107 | potassium ion import | 14.21 | 4 | 1.44E-04 | 3.80E-01 |
| GO:1990573 | potassium ion import across plasma membrane | 14.21 | 4 | 1.44E-04 | 3.17E-01 |
| GO:0055067 | monovalent inorganic cation homeostasis | 6.04 | 7 | 1.46E-04 | 2.74E-01 |
| GO:0015711 | organic anion transport | 3.73 | 11 | 1.66E-04 | 2.73E-01 |
| GO:0055075 | potassium ion homeostasis | 22.65 | 3 | 2.33E-04 | 3.42E-01 |
| GO:0071918 | urea transmembrane transport | 60.4 | 2 | 2.72E-04 | 3.58E-01 |
| GO:0006848 | pyruvate transport | 60.4 | 2 | 2.72E-04 | 3.26E-01 |
| GO:0015840 | urea transport | 60.4 | 2 | 2.72E-04 | 2.99E-01 |
| GO:1901475 | pyruvate transmembrane transport | 60.4 | 2 | 2.72E-04 | 2.76E-01 |
| GO:0045623 | negative regulation of T-helper cell differentiation | 20.13 | 3 | 3.46E-04 | 3.26E-01 |
| GO:0035728 | response to hepatocyte growth factor | 20.13 | 3 | 3.46E-04 | 3.04E-01 |
| GO:0015718 | monocarboxylic acid transport | 6.25 | 6 | 3.66E-04 | 3.01E-01 |
| GO:1902106 | negative regulation of leukocyte differentiation | 7.74 | 5 | 4.21E-04 | 3.26E-01 |
| GO:0034220 | ion transmembrane transport | 2.67 | 15 | 4.51E-04 | 3.30E-01 |
| GO:0072521 | purine-containing compound metabolic process | 3.53 | 10 | 5.14E-04 | 3.56E-01 |
| GO:0043371 | negative regulation of CD4-positive, alpha-beta T cell differentiation | 16.47 | 3 | 6.63E-04 | 4.37E-01 |
| GO:0031167 | rRNA methylation | 16.47 | 3 | 6.63E-04 | 4.16E-01 |
| GO:0009150 | purine ribonucleotide metabolic process | 3.67 | 9 | 7.42E-04 | 4.45E-01 |
| GO:0000466 | maturation of 5.8S rRNA from tricistronic rRNA transcript (SSU-rRNA, 5.8S rRNA, LSU-rRNA) | 40.27 | 2 | 8.07E-04 | 4.62E-01 |
| GO:0016338 | calcium-independent cell-cell adhesion via plasma membrane cell-adhesion molecules | 15.1 | 3 | 8.74E-04 | 4.80E-01 |
| GO:0002822 | regulation of adaptive immune response based on somatic recombination of immune receptors built from immunoglobulin superfamily domains | 5.25 | 6 | 9.34E-04 | 4.92E-01 |
| GO:0098742 | cell-cell adhesion via plasma-membrane adhesion molecules | 4.45 | 7 | 9.53E-04 | 4.83E-01 |

## Discussion

To better understand mechanisms of salinity tolerance in the livebearing fish *Limia perugiae*, we compared patterns of gene expression between a freshwater and a hypersaline population. GO enrichment analysis of differentially expressed genes revealed that the genes upregulated in the saltwater population are largely related to ion transport and maintaining chemical homeostasis, while downregulated genes were associated with processes involved in the cell cycle regulation and protein folding. These results provide insight into how *L. perugiae* has colonized a novel environment and maintains homeostasis under extreme salinity stress. Comparisons of our *Limia* results with pre-existing gene expression data collected from freshwater and saltwater ecotypes of South American silversides (*Odontesthes* spp.: Hughes et al. 2017), three-spine stickleback (*Gasterosteus aculeatus*: Gibbons et al. 2017), and Amur ide (*Leuciscus waleckii*: Xu et al. 2013) indicated that there were few shared differentially expressed genes among all four ecotype pairs. Variation in gene expression was largely shaped by phylogeny rather than environment, and the shared differentially expressed genes among all four pairs showed strong variation in the direction and magnitude of differential expression across lineages. Shared differentially expressed genes were primarily associated with ion transport and immune function. Overall, these results suggest that disparate lineages utilize different mechanisms for overcoming salinity challenges—at least at the level of gene expression. We found that various patterns of gene expression can emerge from crossing saltwater-freshwater boundaries, providing evidence of diverse responses of teleost lineages to a similar environmental challenge. A major question remaining is to what degree variation in gene expression in *L. perugiae* and the other lineages here shaped by phenotypic plasticity and by genetic differences in gene regulation. A previous study in stickleback found evidence for heritable gene expression differences between freshwater and saltwater ecotypes, evidence for shared plastic responses between ecotypes, but only little evidence for ecotype-specific plasticity (Gibbons et al. 2017). Similar studies that combine field studies with laboratory experimentation are highly warranted for other study systems.

### Responses to variation in salinity in a freshwater and hypersaline population of L. perugiae

#### Regulation of transmembrane transport and gill epithelial permeability

Overcoming the physiological challenges associated with transitioning between saltwater and freshwater environments requires modification of ion transport across the gill epithelia (Foskett et al. 1983). In our analysis of differentially expressed genes between a freshwater and hypersaline population of *L. perugiae*, we found evidence for differential regulation of ion transport. Among the genes that were upregulated in the hypersaline population, GO enrichment analysis revealed that several terms were associated with anion transport, sodium ion transport, metal ion transport, and maintaining chemical homeostasis. Genes corresponding to solute carrier families (e.g., *SLC9A3*, *SLC8B1*, *SLC12A8*, *SLC30A9*, *SLC7A1*) and ATPases (e.g., *ATP1B1*, *ATP6V1A*, *ATP13A3*) were among the upregulated genes that are important in maintaining ion and chemical homeostasis (Li et al. 2020). Increasing the rate of inorganic ion, amino acid, and nucleotide transport via upregulation of solute carriers allows aquatic organisms to maintain osmotic balance in saline environments, potentially facilitating acclimation or adaptation to hypersalinity (Li et al. 2020). Specifically, ATPases—such as Na^+^/K^+^-ATPase—have been well-studied for their role in salinity tolerance during both acclimation and adaptation (Deane & Woo 2004; Gonzalez et al. 2005; Langdon & Thorpe 1984; Laverty & Skadhauge 2012; Scott et al. 2004; Tipsmark et al. 2002; Weaver, Tello, et al. 2016). Consistent with our study, previous Western blot analyses of freshwater and hypersaline *L. perugiae* populations have revealed an increase in Na^+^/K^+^-ATPase expression in hypersaline *L. perugiae* when compared to their freshwater conspecifics, which is essential for excreting Na^+^ and Cl^-^ out of the body to maintain homeostasis (Weaver, Tello, et al. 2016).

At very high salinity levels, it is especially difficult to pump Na^+^ against the chemical gradient and out of the gill (Gonzalez 2012). In a hypersaline environment, it therefore may be beneficial to maintain a lower epithelial salt permeability to avoid a back-flux of salts across the tight junctions of the gill epithelia (Gonzalez 2012; Lam et al. 2014). We found up-regulation of genes involved in cell-cell junction organization, including claudins involved in the formation of tight junctions (*CLDN3*, *CLDN4*, *CLDN5*, *CLDN8*, and *CLDN10*). Expression of gill claudin genes has been associated with salinity acclimation in fish (Lam et al. 2014; Tipsmark et al. 2002), and up-regulation of claudin-8 (*CLDN8*) has been shown to reduce the paracellular barrier permeability to Na^+^ (Amasheh et al. 2009), suggesting a role for regulation of gill epithelial permeability via tight junctions during osmoregulatory challenges. Our findings support the hypothesis that modifications to transmembrane transport and gill epithelial permeability jointly contribute to the salt exclusion necessary for survival in a hypersaline environment.

#### Cell cycle regulation

Populations living in hypersaline environments must cope with the adverse effects of high salinity levels, often resulting in strategies for damage repair and energy reallocation (Laverty & Skadhauge 2012; Weaver, Tello, et al. 2016). Functional analysis of the downregulated genes in the hypersaline population of *L. perugiae* showed signatures of downregulation of cell cycle and protein folding processes.

Downregulation of genes involved in the cell cycle is expected to occur under high salinity stress to stop the replication of damaged cells and allow time for DNA repair (Evans & Kültz 2020; Kaufmann & Paules 1996). Similarly, locally adapted killifish living in freshwater and brackish environments exhibit evidence for divergence in the expression of genes underlying control of the cell cycle, supporting a central role in cell cycle regulation in adaptation to salinity stress (Brennan et al. 2015; Evans & Kültz 2020). However, our findings may also suggest a general re-allocation of energy resources (Evans & Kültz 2020). In hypersaline *L. perugiae*, the downregulation of genes involved in the cell cycle and protein folding could be reflective of an energetic trade-off between increased demand for processes involved in osmoregulation and the energetic cost of growth under stressful conditions (Weaver, Tello, et al. 2016). In support of such energetic trade-offs, hypersaline *L. perugiae* have morphological differences from their freshwater counterparts, including a significantly smaller body size and reduced secondary sex features in males (Weaver, Tello, et al. 2016; Haney & Walsh 2003).

#### Regulation of cell signaling

To respond to osmoregulatory challenges, fishes must perceive their environmental salinity and maintain intracellular signals that modulate ion transport and other processes (Evans 2002; Kültz 2012). We found differentially expressed genes associated with a variety of cell signaling processes. Several upregulated genes in the hypersaline population were enriched in GO terms associated with cell signaling, including signal transduction, regulation of signaling, G-protein-coupled receptor signaling pathway, and second-messenger-mediated signaling. G-protein-coupled receptor signaling is among the pathways involved in allowing aquatic organisms to sense their environmental salinity (Papakostas et al. 2012; Sun et al. 2020). Some of these signaling pathway components may also play a role in regulating ion transport, such as mitogen-activated protein kinases (MAPKs) and other serine/threonine protein kinases (Evans 2010; Herzig & Neumann 2000; Kozak et al. 2014; Kültz & Avila 2001), which were among the upregulated genes. MAPK signaling pathways are also implicated in regulation of the cell cycle, some of which trigger cellular growth arrest and DNA damage repair in response to osmotic stress (Kültz & Avila 2001; Kültz & Burg 1998). The upregulation of signaling pathways, especially MAPK genes, may consequently play a role in the downregulation of cell-cycle genes that we found in the hypersaline *L. perugiae* population.

### Comparisons among disparate lineages

#### Lineage-specific responses to variation in salinity

Convergence in gene expression among independently evolved populations can occur in response to shared environmental stressors (Fisher & Oleksiak 2007; Greenway et al. 2020). Among fishes, cases of convergence in gene expression have been identified in response to selective pressures such as pollution (Fisher & Oleksiak 2007), hydrogen sulfide (Greenway et al. 2020), and the absence of light in caves (Stahl & Gross 2017). In contrast, our comparative transcriptomics analysis indicated there is little evidence for convergence in gene expression patterns among fish lineages in response to variation in salinity. Even among the shared differentially expressed genes, there is considerable variation in the magnitude and direction of expression differences between ecotypes across lineages. This finding mirrors genomic analyses that looked for convergent signatures associated with adaptation to salinity variation, which found osmoregulation genes to be common targets of selection but also variation in the exact genes that were involved across different lineages (Velotta et al. 2022).

There are several non-mutually exclusive hypotheses for why we might not expect convergence in gene expression in response to variation in salinity (Kaeuffer et al. 2012). First, changes in both protein-coding DNA sequences and gene expression may occur during adaptation to a novel environment, or they may occur independently (Brown et al. 2019; Jones et al. 2012; Rosenblum et al. 2004). Under certain conditions, selection may favor changes in protein structure and function via modifications at the sequence level, regardless of whether or not gene expression is affected (Brown et al. 2019). Alternatively, changes in gene expression may be favored without selection acting at the sequence level (Brown et al. 2019). Variation in how selection acts on gene expression could contribute to the lack of convergence among the lineages in our study, and there also may be stronger signals of convergence at levels other than gene expression.

Secondly, differences in genetic architecture among lineages may cause different responses to selection, leading to diverse evolutionary outcomes (Kaeuffer et al. 2012). Convergence is therefore less likely to occur with increasing genetic divergence between lineages (Conte et al. 2012), so it is not necessarily surprising that we did not find convergence among distantly related fishes. Even if the genetic architectures are similar among lineages, idiosyncratic responses to selection can also arise as a consequence of functional redundancy (Láruson et al. 2020). Specifically, modification of different genes and pathways may have equivalent functional and fitness consequences (Kozak et al. 2014; Velotta et al. 2022). Functional redundancy is particularly common in complex traits like salinity tolerance that involve many genes and physiological pathways (Láruson et al. 2020). While salinity tolerance is a shared outcome among the lineages in our analysis, the molecular mechanisms underlying osmoregulation may be unique to each lineage.

Third, the directionality of habitat transition may impact gene expression responses, especially if there are genetic adaptations to the habitat of origin. For example, a high-activity version of an ion transporter may be downregulated during transitions to freshwater, while a low-activity version of the same enzyme may be upregulated in the opposite direction. Among the four lineages we included in our study, two of them transitioned from saltwater to freshwater environments (*Gasterosteus* and *Odontesthes*), and two transitioned from freshwater to saltwater environments (*Limia* and *Leuciscus*). If the direction of transition elicits similar responses, then we would expect populations that transitioned in the same direction to share more differentially expressed genes than those that did not transition in the same direction. However, we did not find this to be the case, as the pairs transitioning in the same direction share fewer differentially expressed genes than those that made opposite transitions (Figure 3C).

Finally, covariation with other sources of selection—both abiotic and biotic—may cause idiosyncratic gene expression patterns across lineages. Beyond the challenges directly imposed by variation in salinity, such habitat transitions are often accompanied by other environmental challenges, such as variation in temperature, exposure to novel parasites, and restructuring of host-associated microbial communities (Hughes et al. 2017; Lee & Bell 1999; Lokesh & Kiron 2016; Schmidt et al. 2015). For example, hypersaline Amur ide must cope with high alkalinity stress in addition to salinity, and hypersaline *L. perugiae* have a warmer environment than freshwater *L. perugiae* (Weaver, Tello, et al. 2016; Xu et al. 2013). Additionally, there are differences in the gill microbial communities of saline and freshwater populations of South American silversides (Hughes et al. 2017). Such correlated environmental factors can contribute variation in gene expression responses to salinity transitions we observed.

#### Shared responses involving ion transport and immune function

Although there is little evidence for convergence among all four lineage pairs included in our analysis, there was some functional overlap in the differentially expressed genes shared among three or more lineages. Most of the shared differentially expressed genes were associated with ion transport, as well as some immune system processes. As previously discussed, regulation of ion transport is expected to be crucial when crossing a saltwater-freshwater boundary (Foskett et al. 1983) and osmoregulation genes also exhibit evidence of convergent evolution during salinity transitions (Velotta et al. 2022). Variation in expression of immune genes, particularly those related to inflammation and adaptive immunity, has also been documented in fish during salinity acclimation (El-Leithy et al. 2019; Jeffries et al. 2019). Under a variety of selection pressures, locally adapted populations also frequently show divergence in immune genes due to other factors such as life history differences, different parasite exposure, and shifts in the microbiome (Eizaguirre et al. 2012; Hughes et al. 2017; Jeffries et al. 2019; Zhang et al. 2015; Miller et al. 2001). Immune loci are consequently evolutionary hotspots in diversification (Holmes 2004; Hughes 2002; Sommer 2005). It needs to be tested whether changes in the expression of immune-related genes are directly linked to variation in salinity or whether these genes generally respond to changes in correlated biotic sources of selection.

Overall, our analysis of gene expression patterns between locally adapted freshwater and hypersaline populations of *L. perugiae* revealed upregulation of genes involved in ion regulation, maintaining chemical homeostasis, and cell signaling in the hypersaline population, as well as downregulation of genes involved in the cell cycle and protein folding. These results provide insight into how this livebearing fish maintains homeostasis in a hypersaline environment. Several other lineages of teleosts have similarly made transitions between freshwater and saline environments, providing an opportunity to test the predictability of responses to variation in salinity. We therefore investigated whether disparate lineages show evidence of convergence in gene expression in response to saltwater-freshwater transitions. Our comparisons between four population pairs of freshwater and saline ecotypes in disparate teleost lineages showed little evidence for convergence, as there were only ten differentially expressed genes that were shared among them all. Despite this, we found that the differentially expressed genes shared in three or more of the lineages reflected biological processes related to ion transport and immune functioning.

## Acknowledgements

We thank Ingo Schlupp for help with the field collections and the constructive feedback on the manuscript. Funding was provided by National Science Foundation (IOS-1931657, IOS-2311366), the Army Research Office (W911NF-15-1-0175, W911NF-16-1-0225), and the Des Lee Collaborative Vision in Zoological Studies. We thank the authors of past salinity transcriptomics papers (Hughes et al. 2017; Gibbons et al. 2017; Xu et al. 2013) who made their data publicly available and enabled the comparative analyses presented here.

## Data availability

The data underlying this article are publically available (Table 1).

## References

Amasheh S et al. 2009. Na^+^ absorption defends from paracellular back-leakage by claudin-8 upregulation. Biochem. Biophys. Res. Commun. 378:45–50. doi: 10.1016/j.bbrc.2008.10.164.

Benjamini Y, Hochberg Y. 1995. Controlling the false discovery rate: A practical and powerful approach to multiple testing. J. R. Stat. Soc. 57:289–300. doi: 10.1111/j.2517-6161.1995.tb02031.x.

Betancur-R R. 2010. Molecular phylogenetics supports multiple evolutionary transitions from marine to freshwater habitats in ariid catfishes. Mol. Phylogenet. Evol. 55:249–258. doi: 10.1016/j.ympev.2009.12.018.

Betancur-R R, Ortí G, Pyron RA. 2015. Fossil-based comparative analyses reveal ancient marine ancestry erased by extinction in ray-finned fishes. Ecol. Lett. 18:441–450. doi: 10.1111/ele.12423.

Bloom DD, Lovejoy NR. 2017. On the origins of marine-derived freshwater fishes in South America. J. Biogeogr. 44:1927–1938. doi: 10.1111/jbi.12954.

Bloom DD, Lovejoy NR. 2014. The evolutionary origins of diadromy inferred from a time-calibrated phylogeny for Clupeiformes (herring and allies). Proc. Biol. Sci. 281:20132081. doi: 10.1098/rspb.2013.2081.

Bloom DD, Weir JT, Piller KR, Lovejoy NR. 2013. Do freshwater fishes diversify faster than marine fishes? A test using state-dependent diversification analyses and molecular phylogenetics of new world silversides (Atherinopsidae). Evolution. 67:2040–2057. doi: 10.1111/evo.12074.

Brennan RS, Galvez F, Whitehead A. 2015. Reciprocal osmotic challenges reveal mechanisms of divergence in phenotypic plasticity in the killifish *Fundulus heteroclitus*. J. Exp. Biol. 218:1212– 1222. doi: 10.1242/jeb.110445.

Brown AP et al. 2019. Local ancestry analysis reveals genomic convergence in extremophile fishes. Philos. Trans. R. Soc. Lond. B Biol. Sci. 374:20180240. doi: 10.1098/rstb.2018.0240.

Camacho C et al. 2009. BLAST+: architecture and applications. BMC Bioinformatics. 10:421. doi: 10.1186/1471-2105-10-421.

Carrete Vega G, Wiens JJ. 2012. Why are there so few fish in the sea? Proc. Biol. Sci. 279:2323–2329. doi: 10.1098/rspb.2012.0075.

Chervinski J. 1984. Salinity tolerance of the guppy, *Poecilia reticulata* Peters. J. Fish Biol. 24:449–452. doi: 10.1111/j.1095-8649.1984.tb04815.x.

Coffin JL, Kelley JL, Jeyasingh PD, Tobler M. 2022. Impacts of heavy metal pollution on the ionomes and transcriptomes of Western mosquitofish (*Gambusia affinis*). Mol. Ecol. 31:1527–1542. doi: 10.1111/mec.16342.

Conte GL, Arnegard ME, Peichel CL, Schluter D. 2012. The probability of genetic parallelism and convergence in natural populations. Proceedings of the Royal Society B. 279:5039–5047. doi: 10.1098/rspb.2012.2146.

Corush JB. 2019. Evolutionary patterns of diadromy in fishes: more than a transitional state between marine and freshwater. BMC Evol. Biol. 19:168. doi: 10.1186/s12862-019-1492-2.

Davis AM, Unmack PJ, Pusey BJ, Johnson JB, Pearson RG. 2012. Marine-freshwater transitions are associated with the evolution of dietary diversification in terapontid grunters (Teleostei: Terapontidae). J. Evol. Biol. 25:1163–1179. doi: 10.1111/j.1420-9101.2012.02504.x.

Deane EE, Woo NYS. 2004. Differential gene expression associated with euryhalinity in sea bream (*Sparus sarba*). Am. J. Physiol. Regul. Integr. Comp. Physiol. 287:R1054–63. doi: 10.1152/ajpregu.00347.2004.

Eden E, Navon R, Steinfeld I, Lipson D, Yakhini Z. 2009. GOrilla: a tool for discovery and visualization of enriched GO terms in ranked gene lists. BMC Bioinformatics. 10:48. doi: 10.1186/1471-2105-10-48.

Eizaguirre C, Lenz TL, Kalbe M, Milinski M. 2012. Divergent selection on locally adapted major histocompatibility complex immune genes experimentally proven in the field. Ecol. Lett. 15:723– 731. doi: 10.1111/j.1461-0248.2012.01791.x.

El-Leithy AAA et al. 2019. Optimum salinity for Nile tilapia (*Oreochromis niloticus*) growth and mRNA transcripts of ion-regulation, inflammatory, stress- and immune-related genes. Fish Physiol. Biochem. 45:1217–1232. doi: 10.1007/s10695-019-00640-7.

Emms DM, Kelly S. 2019. OrthoFinder: phylogenetic orthology inference for comparative genomics. Genome Biol. 20:238. doi: 10.1186/s13059-019-1832-y.

Emms DM, Kelly S. 2015. OrthoFinder: solving fundamental biases in whole genome comparisons dramatically improves orthogroup inference accuracy. Genome Biol. 16:157. doi: 10.1186/s13059-015-0721-2.

Erbelding-Denk C et al. 1994. Male polymorphism in *Limia perugiae* (Pisces: Poeciliidae). Behav. Genet. 24:95–101. doi: 10.1007/BF01067933.

Evans DH. 2002. Cell signaling and ion transport across the fish gill epithelium. J. Exp. Zool. 293:336–347. doi: 10.1002/jez.10128.

Evans TG. 2010. Co-ordination of osmotic stress responses through osmosensing and signal transduction events in fishes. J. Fish Biol. 76:1903–1925. doi: 10.1111/j.1095-8649.2010.02590.x.

Evans TG, Kültz D. 2020. The cellular stress response in fish exposed to salinity fluctuations. J Exp Zool A Ecol Integr Physiol. 333:421–435. doi: 10.1002/jez.2350.

Fisher MA, Oleksiak MF. 2007. Convergence and divergence in gene expression among natural populations exposed to pollution. BMC Genomics. 8:108. doi: 10.1186/1471-2164-8-108.

Foskett JK, Bern HA, Machen TE, Conner M. 1983. Chloride cells and the hormonal control of teleost fish osmoregulation. J. Exp. Biol. 106:255–281. doi: 10.1242/jeb.106.1.255.

Gibbons TC, Metzger DCH, Healy TM, Schulte PM. 2017. Gene expression plasticity in response to salinity acclimation in threespine stickleback ecotypes from different salinity habitats. Mol. Ecol. 26:2711–2725. doi: 10.1111/mec.14065.

Gonzalez RJ. 2012. The physiology of hyper-salinity tolerance in teleost fish: a review. J. Comp. Physiol. B. 182:321–329. doi: 10.1007/s00360-011-0624-9.

Gonzalez RJ, Cooper J, Head D. 2005. Physiological responses to hyper-saline waters in sailfin mollies (*Poecilia latipinna*). Comp. Biochem. Physiol. A Mol. Integr. Physiol. 142:397–403. doi: 10.1016/j.cbpa.2005.08.008.

Greenway R et al. 2020. Convergent evolution of conserved mitochondrial pathways underlies repeated adaptation to extreme environments. Proc. Natl. Acad. Sci. U. S. A. 117:16424–16430. doi: 10.1073/pnas.2004223117.

Greenwell MG, Sherrill J, Clayton LA. 2003. Osmoregulation in fish. Mechanisms and clinical implications. Vet. Clin. North Am. Exot. Anim. Pract. 6:169–89, vii. doi: 10.1016/s1094-9194(02)00021-x.

Haney DC, Walsh SJ. 2003. Influence of salinity and temperature on the physiology of *Limia melanonotata* (Cyprinodontiformes : Poeciliidae): a search for abiotic factors limiting insular distribution in Hispaniola. Caribb. J. Sci. 39:327–337.

Harris MA et al. 2004. The Gene Ontology (GO) database and informatics resource. Nucleic Acids Res. 32:D258–61. doi: 10.1093/nar/gkh036.

Herzig S, Neumann J. 2000. Effects of serine/threonine protein phosphatases on ion channels in excitable membranes. Physiol. Rev. 80:173–210. doi: 10.1152/physrev.2000.80.1.173.

Holmes EC. 2004. Adaptation and immunity. PLoS Biol. 2:E307. doi: 10.1371/journal.pbio.0020307.

Hughes AL. 2002. Natural selection and the diversification of vertebrate immune effectors. Immunol. Rev. 190:161–168. doi: 10.1034/j.1600-065x.2002.19012.x.

Hughes LC et al. 2017. Transcriptomic differentiation underlying marine-to-freshwater transitions in the South American silversides *Odontesthes argentinensis* and *O. bonariensis* (Atheriniformes). Ecol. Evol. 7:5258–5268. doi: 10.1002/ece3.3133.

Hwang PP. 1987. Tolerance and ultrastructural responses of branchial chloride cells to salinity changes in the euryhaline teleost *Oreochromis mossambicus*. Mar. Biol. 94:643–649. doi: 10.1007/BF00431411.

Jeffries KM et al. 2019. Divergent transcriptomic signatures in response to salinity exposure in two populations of an estuarine fish. Evol. Appl. 12:1212–1226. doi: 10.1111/eva.12799.

Jones FC et al. 2012. The genomic basis of adaptive evolution in threespine sticklebacks. Nature. 484:55–61. doi: 10.1038/nature10944.

Kaeuffer R, Peichel CL, Bolnick DI, Hendry AP. 2012. Parallel and nonparallel aspects of ecological, phenotypic, and genetic divergence across replicate population pairs of lake and stream stickleback. Evolution. 66:402–418. doi: 10.1111/j.1558-5646.2011.01440.x.

Kaufmann WK, Paules RS. 1996. DNA damage and cell cycle checkpoints. FASEB J. 10:238–247. doi: 10.1096/fasebj.10.2.8641557.

Kelley JL et al. 2012. Genomic resources for a model in adaptation and speciation research: characterization of the Poecilia mexicana transcriptome. BMC Genomics. 13:652. doi: 10.1186/1471-2164-13-652.

Kelley JL et al. 2016. Mechanisms underlying adaptation to life in hydrogen sulfide-rich environments. Mol. Biol. Evol. 33:1419–1434. doi: 10.1093/molbev/msw020.

Kozak GM, Brennan RS, Berdan EL, Fuller RC, Whitehead A. 2014. Functional and population genomic divergence within and between two species of killifish adapted to different osmotic niches. Evolution. 68:63–80. doi: 10.1111/evo.12265.

Kültz D. 2015. Physiological mechanisms used by fish to cope with salinity stress. J. Exp. Biol. 218:1907–1914. doi: 10.1242/jeb.118695.

Kültz D. 2012. The combinatorial nature of osmosensing in fishes. Physiology. 27:259–275. doi: 10.1152/physiol.00014.2012.

Kültz D, Avila K. 2001. Mitogen-activated protein kinases are in vivo transducers of osmosensory signals in fish gill cells. Comp. Biochem. Physiol. B Biochem. Mol. Biol. 129:821–829. doi: 10.1016/s1096-4959(01)00395-5.

Kültz D, Burg M. 1998. Evolution of osmotic stress signaling via MAP kinase cascades. J. Exp. Biol. 201:3015–3021. doi: 10.1242/jeb.201.22.3015.

Kusakabe M et al. 2017. Genetic basis for variation in salinity tolerance between stickleback ecotypes. Mol. Ecol. 26:304–319. doi: 10.1111/mec.13875.

Lam SH et al. 2014. Differential transcriptomic analyses revealed genes and signaling pathways involved in iono-osmoregulation and cellular remodeling in the gills of euryhaline Mozambique tilapia, *Oreochromis mossambicus*. BMC Genomics. 15:921. doi: 10.1186/1471-2164-15-921.

Langdon JS, Thorpe JE. 1984. Response of the gill Na^+^-K^+^-ATPase activity, succinic dehydrogenase activity and chloride cells to saltwater adaptation in Atlantic salmon, *Salmo salar* L., parr and smolt. J. Fish Biol. 24:323–331. doi: 10.1111/j.1095-8649.1984.tb04803.x.

Langfelder P, Horvath S. 2012. Fast R Functions for Robust Correlations and Hierarchical Clustering. J. Stat. Softw. 46. https://www.ncbi.nlm.nih.gov/pubmed/23050260.

Langfelder P, Horvath S. 2008. WGCNA: an R package for weighted correlation network analysis. BMC Bioinformatics. 9:559. doi: 10.1186/1471-2105-9-559.

Langfelder P, Zhang B, Horvath S. 2008. Defining clusters from a hierarchical cluster tree: the Dynamic Tree Cut package for R. Bioinformatics. 24:719–720. doi: 10.1093/bioinformatics/btm563.

Láruson ÁJ, Yeaman S, Lotterhos KE. 2020. The importance of genetic redundancy in evolution. Trends Ecol. Evol. 35:809–822. doi: 10.1016/j.tree.2020.04.009.

Laverty G, Skadhauge E. 2012. Adaptation of teleosts to very high salinity. Comp. Biochem. Physiol. A Mol. Integr. Physiol. 163:1–6. doi: 10.1016/j.cbpa.2012.05.203.

Lee CE, Bell MA. 1999. Causes and consequences of recent freshwater invasions by saltwater animals. Trends Ecol. Evol. 14:284–288. doi: 10.1016/S0169-5347(99)01596-7.

Lee CE, Remfert JL, Gelembiuk GW. 2003. Evolution of physiological tolerance and performance during freshwater invasions. Integr. Comp. Biol. 43:439–449. doi: 10.1093/icb/43.3.439.

Li C et al. 2020. Analysis of differential gene expression in *Litopenaeus vannamei* under high salinity stress. Aquaculture Reports. 18:100423. doi: 10.1016/j.aqrep.2020.100423.

Li H, Durbin R. 2009. Fast and accurate short read alignment with Burrows-Wheeler transform. Bioinformatics. 25:1754–1760. doi: 10.1093/bioinformatics/btp324.

Lokesh J, Kiron V. 2016. Transition from freshwater to seawater reshapes the skin-associated microbiota of Atlantic salmon. Sci. Rep. 6:19707. doi: 10.1038/srep19707.

Love MI, Huber W, Anders S. 2014. Moderated estimation of fold change and dispersion for RNA-seq data with DESeq2. Genome Biol. 15:550. doi: 10.1186/s13059-014-0550-8.

Miller KM, Kaukinen KH, Beacham TD, Withler RE. 2001. Geographic heterogeneity in natural selection on an MHC locus in sockeye salmon. Genetica. 111:237–257. doi: 10.1023/a:1013716020351.

Myers GS. 1949. Salt-tolerance of fresh-water fish groups in relation to zoogeographical problems. Bijdr. Dierkd. 28:315–322. doi: 10.1163/26660644-02801038.

Papakostas S et al. 2012. A proteomics approach reveals divergent molecular responses to salinity in populations of European whitefish (*Coregonus lavaretus*). Mol. Ecol. 21:3516–3530. doi: 10.1111/j.1365-294X.2012.05553.x.

Passow CN et al. 2017. The roles of plasticity and evolutionary change in shaping gene expression variation in natural populations of extremophile fish. Mol. Ecol. 26:6384–6399. doi: 10.1111/mec.14360.

Pertea M et al. 2015. StringTie enables improved reconstruction of a transcriptome from RNA-seq reads. Nat. Biotechnol. 33:290–295. doi: 10.1038/nbt.3122.

Pertea M, Kim D, Pertea GM, Leek JT, Salzberg SL. 2016. Transcript-level expression analysis of RNA-seq experiments with HISAT, StringTie and Ballgown. Nat. Protoc. 11:1650–1667. doi: 10.1038/nprot.2016.095.

Rauchenberger M. 1988. Historical biogeography of poeciliid fishes in the Caribbean. Syst. Zool. 37:356–365. doi: 10.2307/2992198.

Robinson MD, McCarthy DJ, Smyth GK. 2010. edgeR: a Bioconductor package for differential expression analysis of digital gene expression data. Bioinformatics. 26:139–140. doi: 10.1093/bioinformatics/btp616.

Rosen DE, Bailey RM. 1963. The poeciliid fishes (Cyprinodontiformes), their structure, zoogeography and systematics. Bull. Am. Mus. Nat. Hist. 126:1–176.

Rosenblum EB, Hoekstra HE, Nachman MW. 2004. Adaptive reptile color variation and the evolution of the Mc1r gene. Evolution. 58:1794–1808. doi: 10.1111/j.0014-3820.2004.tb00462.x.

Schmidt VT, Smith KF, Melvin DW, Amaral-Zettler LA. 2015. Community assembly of a euryhaline fish microbiome during salinity acclimation. Mol. Ecol. 24:2537–2550. doi: 10.1111/mec.13177.

Scott GR, Richards JG, Forbush B, Isenring P, Schulte PM. 2004. Changes in gene expression in gills of the euryhaline killifish *Fundulus heteroclitus* after abrupt salinity transfer. Am. J. Physiol. Cell Physiol. 287:C300–9. doi: 10.1152/ajpcell.00054.2004.

Smith SA, Bermingham E. 2005. The biogeography of lower Mesoamerican freshwater fishes. J. Biogeogr. 32:1835–1854. doi: 10.1111/j.1365-2699.2005.01317.x.

Sommer S. 2005. The importance of immune gene variability (MHC) in evolutionary ecology and conservation. Front. Zool. 2:16. doi: 10.1186/1742-9994-2-16.

Stahl BA, Gross JB. 2017. A comparative transcriptomic analysis of development in two *Astyanax* cavefish populations. J. Exp. Zool. B Mol. Dev. Evol. 328:515–532. doi: 10.1002/jez.b.22749.

Sun S, Zhu M, Pan F, Feng J, Li J. 2020. Identifying neuropeptide and G-protein-coupled receptors of juvenile oriental river prawn (*Macrobrachium nipponense*) in response to salinity acclimation. Front. Endocrinol. 11:623. doi: 10.3389/fendo.2020.00623.

Therkildsen NO, Palumbi SR. 2017. Practical low-coverage genomewide sequencing of hundreds of individually barcoded samples for population and evolutionary genomics in nonmodel species. Mol. Ecol. Resour. 17:194–208. doi: 10.1111/1755-0998.12593.

Tipsmark CK et al. 2002. Dynamics of Na^+^K^+^,2Cl^-^ cotransporter and Na^+^,K^+^-ATPase expression in the branchial epithelium of brown trout (*Salmo trutta*) and Atlantic salmon (*Salmo salar*). J. Exp. Zool. 293:106–118. doi: 10.1002/jez.10118.

Tsai J-W et al. 2018. A field and laboratory study of the responses of cytoprotection and osmoregulation to salinity stress in mosquitofish (*Gambusia affinis*). Fish Physiol. Biochem. 44:489–502. doi: 10.1007/s10695-017-0448-y.

Velotta JP et al. 2017. Transcriptomic imprints of adaptation to fresh water: parallel evolution of osmoregulatory gene expression in the Alewife. Mol. Ecol. 26:831–848. doi: 10.1111/mec.13983.

Velotta JP, McCormick SD, Whitehead A, Durso CS, Schultz ET. 2022. Repeated Genetic Targets of Natural Selection Underlying Adaptation of Fishes to Changing Salinity. Integr. Comp. Biol. doi: 10.1093/icb/icac072.

Warren WC et al. 2018. Clonal polymorphism and high heterozygosity in the celibate genome of the Amazon molly. Nat Ecol Evol. 2:669–679. doi: 10.1038/s41559-018-0473-y.

Weaver PF, Tello O, et al. 2016. Hypersalinity drives physiological and morphological changes in *Limia perugiae* (Poeciliidae). Biol. Open. 5:1093–1101. doi: 10.1242/bio.017277.

Weaver PF, Cruz A, Johnson S, Dupin J, Weaver KF. 2016. Colonizing the Caribbean: biogeography and evolution of livebearing fishes of the genus *Limia* (Poeciliidae). J. Biogeogr. 43:1808–1819. doi: 10.1111/jbi.12798.

Whitehead A, Roach JL, Zhang S, Galvez F. 2011. Genomic mechanisms of evolved physiological plasticity in killifish distributed along an environmental salinity gradient. Proc. Natl. Acad. Sci. U. S. A. 108:6193–6198. doi: 10.1073/pnas.1017542108.

Xu J et al. 2013. Gene expression changes leading extreme alkaline tolerance in Amur ide (*Leuciscus waleckii*) inhabiting soda lake. BMC Genomics. 14:682. doi: 10.1186/1471-2164-14-682.

Xu P et al. 2014. Genome sequence and genetic diversity of the common carp, *Cyprinus carpio*. Nat. Genet. 46:1212–1219. doi: 10.1038/ng.3098.

Yang W-K, Kang C-K, Chen T-Y, Chang W-B, Lee T-H. 2011. Salinity-dependent expression of the branchial Na^+^/K^+^/2Cl^-^ cotransporter and Na^+^/K^+^-ATPase in the sailfin molly correlates with hypoosmoregulatory endurance. J. Comp. Physiol. B. 181:953–964. doi: 10.1007/s00360-011-0568-0.

Zhang B, Horvath S. 2005. A general framework for weighted gene co-expression network analysis. Stat. Appl. Genet. Mol. Biol. 4:Article17. doi: 10.2202/1544-6115.1128.

Zhang R et al. 2015. Local adaptation of *Gymnocypris przewalskii* (Cyprinidae) on the Tibetan Plateau. Sci. Rep. 5:9780. doi: 10.1038/srep09780.

